# Uncovering Effects of Schizophrenia upon a Maximally Significant, Minimally Complex Subset of Default Mode Network Connectivity Features

**DOI:** 10.1101/2024.04.24.590969

**Authors:** Masoud Seraji, Charles A. Ellis, Mohammad S.E. Sendi, Robyn L. Miller, Vince D. Calhoun

**Author notes:** Funding for this work is provided by NSF grant 2112455 and NIH grant R01EB027147.

## Abstract

A common analysis approach for resting state functional magnetic resonance imaging (rs-fMRI) dynamic functional network connectivity (dFNC) data involves clustering windowed correlation time-series and assigning time windows to clusters (i.e., states) that can be quantified to summarize aspects of the dFNC dynamics. However, those methods can be dominated by a select few features and obscure key dynamics related to less dominant features. This study presents an iterative feature learning approach to identify a maximally significant and minimally complex subset of dFNC features within the default mode network (DMN) in schizophrenia (SZ). Utilizing dFNC data from individuals with SZ and healthy controls (HC), our approach uncovers a subset of features that has a greater number of dFNC states with disorder-related dynamics than is found when all features are present in the clustering. We find that anterior cingulate cortex/posterior cingulate cortex (ACC/PCC) interactions are consistently related to SZ across the most significant iterations of the feature learning analysis and that individuals with SZ tend to spend more time in states with greater intra-ACC anticorrelation and almost no time in a state of high intra-ACC correlation that HCs periodically enter. Our findings highlight the need for nuanced analyses to reveal disorder-related dynamics and advance our understanding of neuropsychiatric disorders.

## I. Introduction

Dynamic functional network connectivity (dFNC) extracted from resting state functional magnetic resonance imaging (rs-fMRI) is a key measure of neural activity that has provided insight into neuropsychiatric disorders [1]–[3] and cognitive function [4], [5]. A common dFNC analysis involves computing correlation within time windows followed by clustering to assign them to clusters (i.e., states) [1]–[3]. The passage of a study participant through those states can then be quantified and provide novel insights. Nevertheless, previous studies of disorders like schizophrenia (SZ) have shown that some dFNC features can dominate clustering analyses and obscure disorder-related dynamics in less-influential subsets of dFNC features [2], [6], [7]. For example, whole-brain analyses [6] have failed to detect disorder-related dynamics found in domain-specific analyses [2]. In this study, we present a novel iterative feature learning approach for uncovering a maximally significant, minimally complex subset of dFNC features (MSMC, i.e., smallest subset of features that also has the most disorder-related dynamics) in the default mode network (DMN) in SZ.

Historically, static functional network connectivity (sFNC; i.e., the correlation between the activation of different brain regions across a recording) has played a key role in rs-fMRI exploration [8], [9]. Nevertheless, multiple studies have shown that sFNC can obscure disease-related dynamics in rs-fMRI data [10], and dFNC (i.e., the correlation of different brain region activations at different points in time), has grown in popularity.

As it has grown in popularity, dFNC has helped characterize many neurological and neuropsychiatric disorders [1]–[3] and aspects of cognition [4], [5]. While many approaches have been developed to analyze dFNC [11], one of the most common approaches involves the use of clustering [1]–[3]. In these analyses, FNC is computed within time windows followed by assignment to clusters. Those clusters are considered to be states, and the passage of a study participant through those states can be summarized using a variety of measures and related to conditions of interest. Common summary measures [12] include the occupancy rate (OCR; i.e., percent of time steps spent in each state) and number of state transitions (NST).

Importantly, as would be expected, some features have a greater impact upon clustering than others and can dominate clustering analyses [7], [13]. The effect of this is that those dominant features are the driving force behind any uncovered dynamics and that other disorder-related dynamics that may be found within a subset of features are obscured. For example, whole-brain analyses [6] have failed to uncover disorder-related dynamics found in domain-specific analyses [2], and domain-specific analyses [2] have failed to uncover disorder-related dynamics within a smaller intra-domain subset of features [7].

As such, in this study, we present an approach for identifying a subset of dFNC features that maximizes the number of significant summary measures associated with the dFNC dynamics. We assign data from the DMN of individuals with SZ (SZs) and healthy controls (HCs) to 5 dFNC states, use an explainable approach [13] to identify key features, remove the feature most important to the clustering, and repeat until the number of states with significant OCRs differentiating SZs and HCs is maximized and the number of included features is minimized. Importantly, our methodology reveals a subset of dynamic functional network connectivity (dFNC) features that not only surpasses the initial clustering in state significance but also consists of a notably reduced number of dFNC features. We find that anterior and posterior cingulate cortex (ACC and PCC) interactions dominate the iterations with greatest significance and that increased intra-ACC anticorrelation in SZs at the MSMC iteration suggest ACC-related dysfunction in SZ.

## II. Methods

Fig. 1 shows an overview of our methodology. We assigned dFNC data from SZs and HCs to five clusters (i.e., states) using k-means clustering with correlation distance. Employing a feature learning approach, we iteratively removed the most significant feature and reclustered. This process continued until the number of features prevented the calculation of correlation distance. We next investigated potential changes in the temporal distribution of the five states across iterations by computing OCRs for each subject and state. We compared OCRs across the SZ and HC groups to identify any differences.

**Fig. 1.**
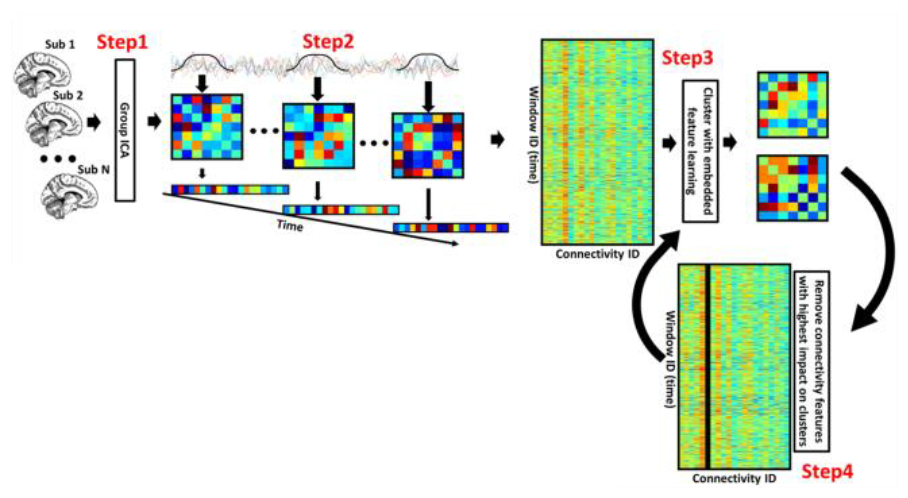
Overview of Methods. In Step 1, we preprocess the data and extract independent components (ICs) from the default mode network (DMN). In **S**tep 2, we compute the correlation between each DMN IC. In **S**teps 3 and 4, the data undergo clustering, incorporating G2PC feature importance to pinpoint key features. The process involves iteratively clustering and removing the most important feature until only a small subset of features remain. This sequence of steps is repeated to systematically distill the relevant information.

### A. Description of Dataset

We employed the Functional Imaging Biomedical Informatics Research Network (FBIRN) dataset, comprising rs-fMRI recordings from 151 SZs and 160 HCs. The dataset has been used in multiple studies [7]. The data were collected across seven sites: the University of California at Irvine, the University of California at Los Angeles, the University of California at San Francisco, Duke University/the University of North Carolina at Chapel Hill, the University of New Mexico, the University of Iowa, and the University of Minnesota. Participants from all sites provided written informed consent through processes approved by local institutional review boards.

### B. Description of Preprocessing

Before preprocessing, we excluded the initial five scans. Subsequently, we applied statistical parametric mapping (SPM12, https://www.fil.ion.ucl.ac.uk/spm/) for preprocessing, incorporating rigid body motion correction to address head motion. Spatial normalization to the echo-planar imaging template in the standard Montreal Neurological Institute space was conducted with resampling to 3x3x3 mm^3^. We applied Gaussian kernel smoothing with a full width at half maximum of 6mm. After preprocessing, independent components (ICs) were extracted using the NeuroMark pipeline [14] in the GIFT toolbox (http://trendscenter.org/software/gift). ICs exhibiting peak activation in the default mode network (DMN) gray matter were deemed relevant to our research. Specifically, we identified seven DMN nodes, including three Precuneus (PCN) nodes, two ACC nodes, and two PCC nodes (Fig. 2).

**Fig. 2.**
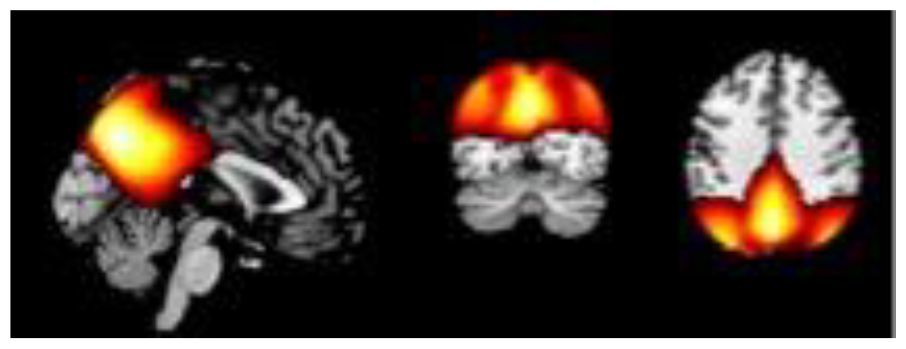
The sagittal, coronal, and axial views, from left to right, of the spatial map of the default mode network.

### C. Description of Feature Learning Approach

To examine the dFNC of the seven DMN nodes of each participant, we used a sliding tapered window with a window size of 20 TRs (40 s). This yielded 21 FNC features (i.e., FNC matrices sized 7x7, corresponding to the seven DMN nodes). Based on previous studies [2], we first assigned samples to five clusters using k-means clustering with correlation distance. Next, to identify the features that most contributed to the clustering, we used Global Permutation Percent Change (G2PC) feature importance [13]. G2PC gauges the sensitivity of clusters to perturbation. It involves permuting a feature across all samples in a dataset and then determining the percentage of samples that change clusters after permutation. Features leading to the highest number of samples switching clusters are deemed the most important, while features that do not prompt such changes are considered less crucial to the clustering process. After identifying the most important feature, we eliminated it and then iteratively repeated the process of clustering the data and removing the key features until two features remained. One of the last two features could not be eliminated because our correlation metric could not be computed with two features. To ensure comparability of states found in each iteration, we performed 500 initializations for the first clustering iteration and initialized later iterations with the cluster centroids (minus the removed feature) of the previous iteration.

### D. Description of MSMC Subset Selection Appoach

Upon identifying dFNC states through our feature learning approach, we investigated potential disparities in the prevalence of these states in SZs and HCs. To quantify this, we computed OCRs for each subject and each of the five states. To evaluate the impact of feature removal on the OCRs of SZs and HCs, we conducted two-sample t-tests comparing SZ and HC OCRs at each iteration. We next identified the MSMC iteration in which differences in the OCRs between HCs and SZs were significant across the largest number of states. While multiple iterations had the same number of significant states, this iteration also had the fewest number of features (minimally complex) with that same number of significant states.

## III. Results AND Discussion

In this section, we describe and discuss the OCR significance and feature removed at each iteration, the MSMC subset of dFNC features, disorder-dependent differences in OCRs, and the limitations and future work of our study.

### A. Identifying Distribution of Significance Across Iterations

As shown in Fig. 3, we calculated the number of states with significant OCRs at each iteration. While we observed 2 significant OCRs in early iterations, after removing around half of the features, the number of significant OCRs peaked at 3 OCRs for 4 iterations. In these four iterations, ACC1/PCN2, ACC1/PCC1, ACC1/PCC2, and ACC1/ACC2 were the most important features. This suggests that while ACC/PCC interactions were not the most influential features when features from earlier iterations were present, they were likely the most impacted by SZ. In further analyses, we analyzed the iteration for which ACC1/ACC2 was most important to the clustering. These findings support our earlier assertion that the dFNC features most influential to the clustering can obscure disorder-related dynamics and that the removal of those features uncovers those obscured dynamics.

**Fig. 3.**
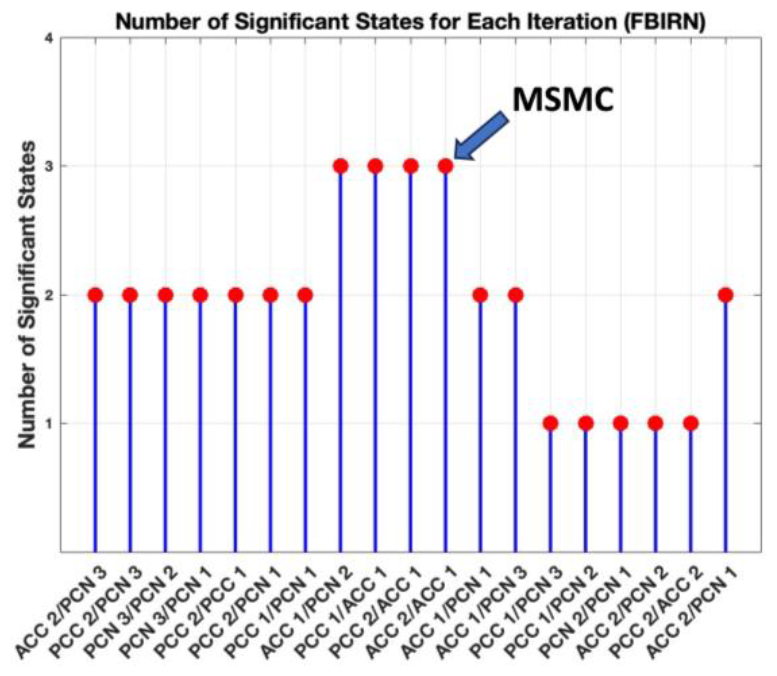
Number of significant states based on the OCR. The x-axes show from left to right the node removed at each iteration, and the y-axis shows the number of significant OCRs. The blue arrow shows the MSMC feature.

### B. Characterizing dFNC States in MSMC Feature Subset

Another analysis we conducted involved identifying an MSMC subset of features. Fig. 4 shows each cluster centroid at the MSMC iteration. States 0 and 2 were characterized by high intra-PCN correlation, while states 1, 4, and 3 had decreasing levels. States 0 and 3 and states 1 and 4 had high and moderate ACC1/PCN1 correlation, respectively, while only state 1 had moderate ACC/PCN3 correlation. State 3 was the only state with low PCC/PCN2 correlation and moderate intra-ACC correlation. In contrast, states 0 and 4 had moderate intra-ACC anticorrelation. States 0 and 2 had moderate PCN/ACC2 anticorrelation, while other states had a mix of PCN/ACC2 correlation and anticorrelation. Lastly, state 4 had a mix of low to moderate PCC/ACC2 correlation and anticorrelation.

**Fig. 4.**
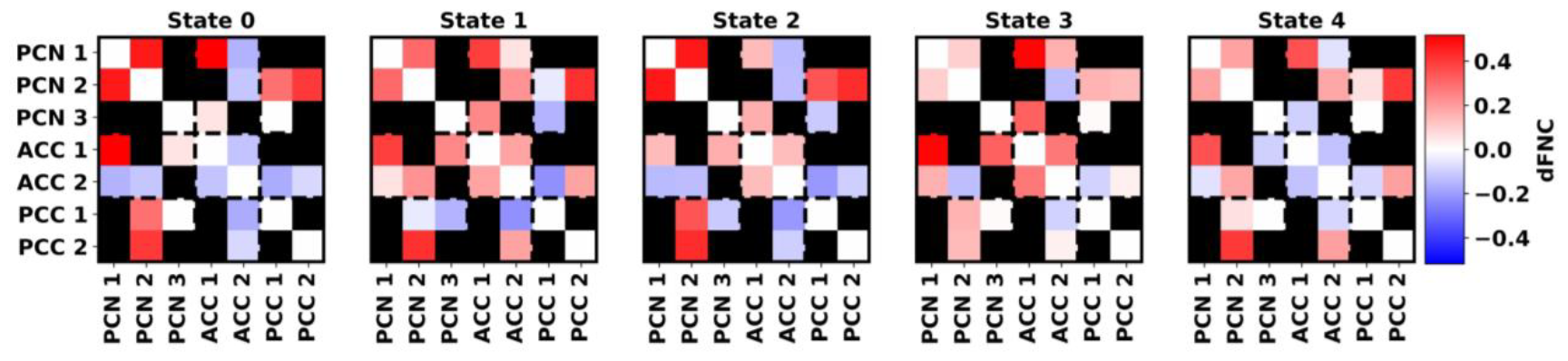
The centroids for the MSMC iteration. Centroids for states 0 through 4 are displayed from left to right. The centroids are displayed as connectivity matrices, showing a heatmap of FNC between each pair of ICs. The corresponding ICs are shown on the x- and y-axes. The color black indicates that the corresponding feature was eliminated. All panels share the same color bar to the right of the figure.

### C. Uncovering Disorder-Dependent Differences in OCRs

Fig. 5 shows the OCR analysis results. Interestingly, significantly different OCRs were observed in state 0, for which both HCs and SZs had the highest OCRs and in states 3 and 4 for which both HCs and SZs had the lowest OCRs, while the states for which they both had moderate OCRs (states 1 and 2) had no significance. SZs spent more time in states 0 and 4 than HCs, suggesting that SZs can be characterized by high PCN/PCC correlation, moderate PCC1/ACC anticorrelation, and moderate intra-ACC anticorrelation. In contrast, HCs favored state 3, the least common state, more than SZs. This suggested that HC activity is marked by moderate to high PCN/PCC correlation, high PCC1/ACC correlation, and moderate intra-ACC correlation. Our PCN/PCC and PCC1/ACC findings fit with those of [12] and [7], respectively.

**Fig. 5.**
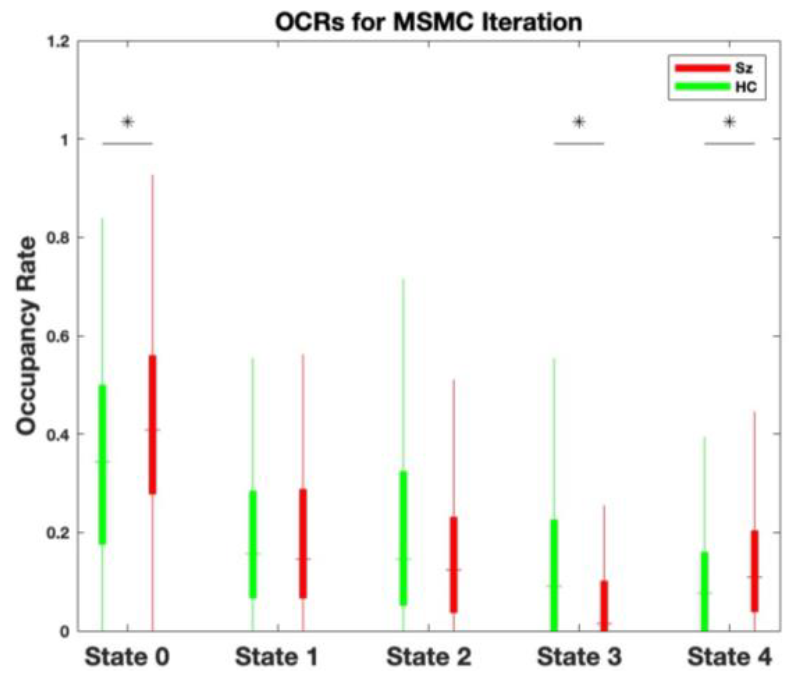
Boxplots of OCRs at the MSMC iteration, (i.e., iteration number 11, removal of ACC2/ACC1). HCs and SZs are shown in green and red, respectively. Asterisks identify differences in SZ and HC OCRs in a state. The x-axis shows each dFNC state, and the y-axis shows the OCR.

Additionally, while both HCs and SZs spend large amounts of time in states with intra-ACC anticorrelation (states 0 and 4) or moderate intra-ACC correlation (states 1 and 2), SZs have more frequent intra-ACC anticorrelation (states 0 and 4), and HCs periodically enter a state of much stronger intra-ACC correlation (state 3) that SZs seldom, if ever, enter. This strongly hints at intra-ACC dysfunction in SZ and builds upon a compelling foundation of existing research [15].

### D. Limitations and Future Work

In this study, we used G2PC as a feature importance metric. While G2PC may not obtain reliable results in high dimensional data spaces, it should theoretically work well for the relatively low number of 21 DMN features we analyzed in this study. Given that samples may be located near the boundaries of their assigned cluster and have similarities to other clusters, future applications of our approach might use fuzzy clustering or meta-state analyses. Also, we only maximized significance for the OCR but could include other summary features (e.g., NST). Lastly, in future studies, it could be helpful to account for the effect size related to significance rather than only the number of significant features.

## IV. Conclusion

Previous studies have suggested that the use of clustering to identify dFNC states may obscure important neuropsychiatric disorder dynamics. Our study introduced a new feature learning approach that involves iteratively identifying and removing key features and then reclustering. Our study focused on the default mode network in schizophrenia and revealed key differences between individuals with schizophrenia and healthy controls after the removal of key features. Importantly, we find that ACC/PCC connectivity was the most disorder-relevant across iterations and that SZs spend more time in states with intra-ACC anticorrelation. This validates our initial hypothesis and promises to advance a more nuanced understanding of neuropsychiatric and neurological disorders in future research.

## V. Acknowledgements

We thank those who collected the FBIRN dataset.

